# Energetic costs of bill heat exchange demonstrate contributions to thermoregulation at high temperatures in toco toucans (*Ramphastos toco*)

**DOI:** 10.1101/2022.11.04.515157

**Authors:** Jussara N. Chaves, Glenn J. Tattersall, Denis V. Andrade

## Abstract

Body temperature regulation in the face of changes in ambient temperature and/or in metabolic heat production involves adjustments in heat exchange rates between the animal and the environment. One of those mechanisms include the modulation of the surface temperature of specific areas of the body through vasomotor adjustment and blood flow control, to change the thermal conductance of this region, thereby promoting dissipation or conservation of body heat. In homeotherms, this thermoregulatory adjustment is essential for the maintenance of body temperature over a moderate temperature range, known as the thermal neutral zone (TNZ), without increasing metabolic rate (MR). Thermal windows are poorly insulated body regions and highly vascularized that are particularly efficient for heat dissipation through that mechanism. The bill of the toco toucan (*Ramphastos toco*) has been described as a highly efficient thermal window and hypothesized to assist in the thermal homeostasis of this bird. Herein, we directly evaluated the contribution of heat exchange through the bill of the toco toucan and role of the bill in the delimitation of the TNZ. To do this, we measured metabolic rate, via oxygen consumption, over a range of ambient temperature from 0 to 35°C (every 5°C). MR measurements were made in birds with the bill intact (control group) and with the bill artificially insulated (experimental group). The limits of the TNZ, 10.9-25.0°C for the control group and 10.8-24.1°C for the experimental group, did not differ between the treatments. MR differed among treatments only at elevated temperatures (30 and 35°C), reaching values of 0.97 ml O_2_·g^-1^·h^-1^·°C^-1^ (± 0.06) for the control group and 1.20 ml O_2_·g^-1^·h^-1^·°C^-1^ (± 0.07) for the experimental group at 35°C. These results indicate that while heat dissipation through the bill does not contribute significantly to widening of the TNZ, it may well be critically important in assisting body temperature regulation at higher temperatures extending above the upper limit of the TNZ. We estimate that the contribution of the bill to total heat exchange approaches 31% of basal metabolic heat production, providing evidence of the substantial role of peripheral heat exchange and linking the role of appendage size as a key factor in the evolution of thermoregulatory responses in endotherms.

## Introduction

Body temperature regulation in endothermic animals involves the balance between the rate of metabolic heat production and the rate of heat exchange between the animal and the environment (Dawson and Whittow, 2000; McNab, 1974). The latter is dictated by the animal’s thermal conductance which, in turn, is influenced by body size and shape, both of which depend on surface/volume relations, body composition, and effectiveness of insulation (Tattersall et al., 2012). Within specific limits of ambient temperatures, body temperature (T_b_) can be maintained reasonably constant by the modulation of thermal conductance, effected by changes in body posture, insulation thickness, or resulting from vasomotor adjustments (Gordon, 2012; IUPS, 2001; Stager et al., 2020). This thermoneutral zone (TNZ) is limited by a lower (LCT) and upper (UCT) critical temperature, beyond which, the regulation of body temperature will require increments in energy expenditure (McNab, 2012). While little controversy exists about LCT, the definition and identification of the UCT is complicated by the fact that it is also defined as the point in which evaporative cooling starts to increase (IUPS, 2001), which often does not coincide with the increment in metabolism, and usually do not appear as an obvious or marked threshold (Gordon, 2012; Long et al., 2014). In any case, the more efficiently an animal can modulate its rate of sensible (dry) heat exchange, the broader should be its TNZ. This is obviously of adaptive value by allowing animals to regulate body temperature at minimum costs (energy or body water) as ambient temperature varies (see Bozinovic et al., 2014).

In birds, a widespread thermoregulatory response contributing to the modulation of sensible heat exchange is the adjustment in the insulative power of the plumage. The feather’s angle of insertion responds to variation in ambient temperature, which alters the thickness of the feather/air insulative layer (Saarela et al., 1984); a mechanism akin to piloerection in mammals (e.g., Hohtola et al., 1980). Also, birds can modulate heat exchange by altering blood perfusion to peripheral organs, such as feet, legs, bill, and head ornaments (Eastick et al., 2019; Hagan and Heath, 1980; Steen and Steen, 1965). These appendages are uninsulated and, as their superficial temperature is altered – via vasomotor adjustments – conductive and radiative heat exchange can be modulated (e.g., Jessen, 2001). In this context, bill size has been demonstrated to vary with latitudinal gradient in environmental temperature (i.e., Allen’s rule; Symonds and Tattersall, 2010), possibly under the influence of the most critical temperatures experienced at different seasons by different species or populations at different geographic locations (see Danner and Greenberg, 2015). Altogether, an adaptive association between appendage (e.g., bill, limbs) size and environmental temperatures and, consequently, to heat exchange capability, seems to be quite well supported (Playa-Montmany et al., 2021; Tattersall et al., 2018). This may involve energetic considerations and we posit that bill size will correlate with the amplitude of the TNZ in birds, particularly in the setting of the UCT threshold. However, the contribution and ability to modulate heat exchange through the bill to keep thermal balance at the expense of minimum metabolic cost, remains untested in birds.

The thermoregulatory role played by the avian bills is epitomized by the toco toucan, whose ostentatious appendage can account for up to 50% of total body surface area and through which this bird has been estimated to dissipate as much heat as 4 times its basal rate of heat production (Tattersall et al., 2009). As such, the toco toucan bill is recognized as the most potent thermal window, through which sensible heat exchange can be adjusted, identified in the animal kingdom. In the present study, we aimed to quantify the contribution of the toco toucan bill in establishing the limits to which body temperature can be maintained at minimum costs. Is the impressive capacity for sensible heat exchange through the bill reflected in the breadth of its TNZ? If that is the case, does having such a large heat exchange organ extend the TNZ’s lower, upper, or both critical thermal limits? We tested these questions by measuring the rates of oxygen uptake 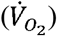 of adults of toco toucans at a range of ambient temperatures under two conditions. First, measurements were made in intact birds, able to use their bill for modulating sensible heat exchange to its full extent. We also artificially insulated the entire bill of the birds effectively disabling the contribution of this heat exchange avenue to the global maintenance of the bird’s thermal balance. By comparing both conditions, we were able to isolate the bill’s contribution to the amplitude and the limits of the TNZ.

## Material and Methods

### Animals

Experiments were carried out with six adults of both sexes of toco toucans, *Ramphastos toco* (Müller, 1776). Birds were obtained by donation from accredited animal facilities in São Paulo state (ICMBio # 27171-1). In captivity, toucans were maintained in outdoor wired mesh cages (6 × 10 m in area, 3 m high) at the Comparative Animal Physiology Laboratory, Universidade Estadual Paulista, Rio Claro municipality, São Paulo state, southeastern Brazil. Cages were provided with shelters, perches, water, and a few small trees; ground surface was mostly covered with grass and other low greens. Toucans were fed with fresh fruits and commercial pelleted food (Labcon Club Toucan Tucanes) offered twice a day.

### Experimental protocol

We measured 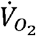 in birds subjected to temperatures varying from 0 to 35ºC, at 5ºC increments. All birds were measured at all experimental temperatures under two conditions: *Intact*, in which birds were measured without any manipulation of the bill’s thermal conductivity, and; *Insulated*, in which the entire external surface of the bill was covered with a layer of heat exchange insulating material. To create this insulation, we used a 2 mm felt cloth tailored to match the exact shape and size of each bird’s bill individually. Felt molds were lined internally with an aluminum foil tape and fixed to the bill with double-sided tape (Figure 1). Thermal conductivity of this heat insulating material was experimentally determined and averaged 0.0729 (±0.006) W·m^-1^·K^-1^ at temperatures varying from 4.3 to 31.5ºC (see Supplementary Material). Upper and lower bill parts were independently covered, so birds could freely engage in gaping and panting during experimentation.

**Figure 1.**
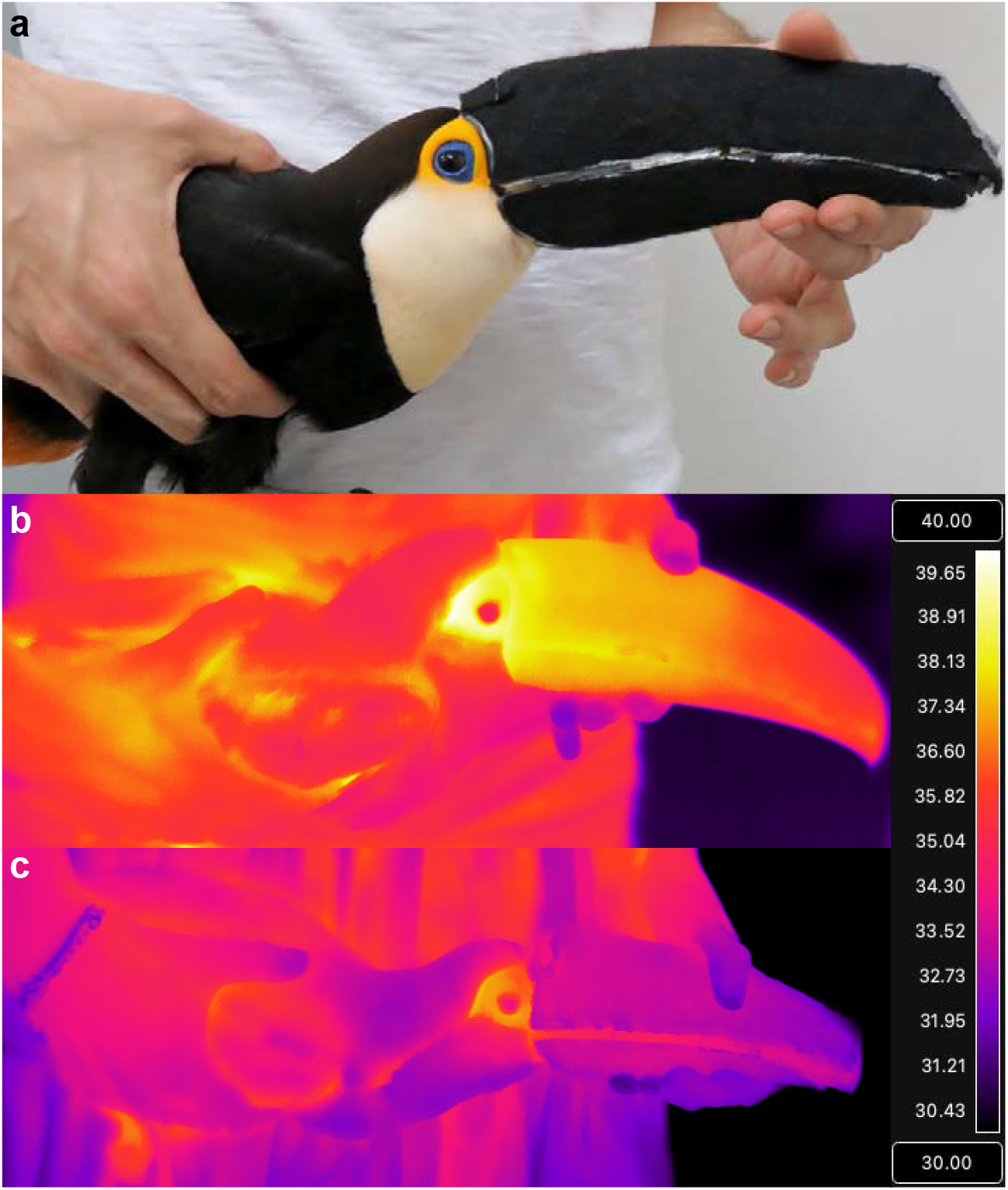
Sample images of insulating material used to reduce heat exchange from the bill of Toco toucans (*Ramphastos toco*). A live image of an insulated bill is depicted in a, while thermal image in b shows a toucan (exposed to 35°C) with an uninsulated bill with higher surface temperature compared to a toucan (exposed to 30°C) with an insulated bill in panel c.

All measurements were taken during the night, which corresponds to the circadian phase when *R. toco* is inactive (Sick and Barruel, 1988). Birds were fasted for 6 to 8 hours previous to the experiments to ensure they were post-absorptive at the time of measurements (Silva et al., 2008). In a typical trial, a fasted bird was transferred from the maintenance cage in the evening, weighed, and placed in a custom-made respirometric chamber (46.5 L, 44 × 44 × 24 cm) housed inside a BOD incubator (FANEM mod. 347 CD) for temperature control. Inside this chamber, we placed a small wood perch, 10 cm from the ground, that the birds invariably sat. The respirometry system was immediately activated, but, in all cases, a habituation period of at least 2 hours was observed before the beginning of data collection. Once started, data collection persisted for approximately 2.5 hours, which proved entirely adequate in yielding consistent steady-state readings. At the end of the experiments, birds were removed from the respirometric chamber, and their core body temperature measured by the insertion of an external temperature probe into their cloaca (Techline, model TS-101).

Individual birds were measured once in each treatment/temperature combination, in random order, and were allowed to recover for at least 3 days between trials. All procedures were approved by the “Animal Use and Ethics Committee” (CEUA) of the Bioscience Institute, São Paulo State University (UNESP), campus of Rio Claro, Brazil (Protocol # 5373).

### Respirometry

We determined the rates of oxygen uptake using an open-flow respirometric system (Lighton, 2008; Voigt and Cruz-Neto, 2009). In our setup, room air was drawn through an in-line column filled with drying agent (Drierite™, W.A. Hammond Drierite CO. LTD) and then ventilated to the animal chamber at a rate of 2.4 L·min^-1^ (MFC 2 -Sable Systems, USA). From the outflow port, we use a custom-made 60 mL syringe manifold to sub-sample the excurrent airflow leaving the respirometric chamber the airstream at a rate of 150 mL^-1^·min^-1^. This sub-sampling line was directed through a drying column (same as above) and then into an oxygen analyzer (FOXBOX; Sable Systems International, USA). Immediately before and after each experimental trial, we also determined the baseline levels of fractional oxygen concentration by running our respirometry system under the identical experimental conditions, but with no animal inside the respirometric chamber.

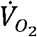 was calculated by the change in fractional oxygen concentration between baseline and that averaged for a steady-state period of at least 15 min measured in the last hour of the bird’s measurement period. Calculations followed Koteja (1996) assuming a Respiratory Exchange Ratio of 0.8.

### Energetic Costs

Heat production (HP; Watts) was estimated from 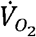 assuming an oxycaloric equivalent of 20 J·mL O_2_^-1^ (Blaxter, 1989), and dividing resulting values by 3600 (to convert h^-1^ to s^-1^), yielding J·s^-1^ or Watts). At each measurement temperature, because experiments were conducted pairwise on the same individuals, we subtracted the control condition HP from the insulated bird’s HP to assess the change, if any, in the energetic costs associated with minimizing bill heat transfer. These values provided an assessment of the energetic cost of unfettered heat exchange across the bill and allowed us to estimate the energetic costs of compensatory mechanisms that might have come into play when the bill heat exchange capacity was disabled. As a result, we were able to estimate the energetic cost of thermoregulation of the heat exchange organ, the bill, provided that insulated birds maintained the same T_b_ as control birds. Finally, for the highest temperature exposure (i.e., above the upper critical temperature), we estimated the hydroregulatory cost for the augmentation in evaporative cooling under the situation in which bill heat transfer was prevented. To this aim, we converted the additional energy expended in the bill insulated group, compared to the control uninsulated one, into the amount of water needed to be evaporated (g H_2_O·day^-1^) to defend body temperature, in doing so, we assumed the heat of vapourisation of water of 2260 J·g^-1^.

### Data Analysis

We calculated thermal conductance at all temperatures using the formula described by McNab (1980):

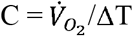

where C is the conductance (ml O_2_·g^-1^·h^-1^·°C^-1^), 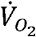 is the metabolic rate (MR; ml O_2_·g^-1^·h^-1^) and ΔT is the difference between body and ambient temperature. Statistical analyses were performed in R version 4.2.0 (R. Developmental (Core) Team, 2020). To examine simultaneously the effects of ambient temperature and insulation treatment, linear mixed-effects models were fit using the package *lme4* (Bates et al., 2015), including the interaction term and animal identify as a random effect, while significance of the models was tested using the *lmerTest* package correcting for degrees of freedom using the Satterthwaite’s method (Kuznetsova et al., 2017). Segmented regressions were used to estimate LCT and UCT from 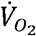 data, and the inflection point in thermal conductance, with respect to ambient temperature, using the *segmented* package (Muggeo, 2017). Plots were generated using the package *ggplot2* (Wickham, 2016).

## Results

The body mass of the toucans calculated at the end of each experiment was slightly lower (by 16.7 grams) in the bill insulated group (619.5 ± 60.6 grams) compared to the intact condition (636.1 ± 65.0 grams; *F*_1,6_=7.93; *P*=0.031), although there was no interaction between treatment and ambient temperature on body mass (*F*_7,90_=1.58; *P*=0.151). The lower and upper limits of the toucan’s TNZ, calculated by manual regression line intersection or according to the three-phase regression (Table 1), did not differ between treatments. The TNZ limits were 10.9 - 25.0°C for the intact group and 10.8 - 24.1°C for the bill insulated group (Table 1). The mass-specific metabolic rate as a function of room temperature was different between treatments due to a greater increase in MR in the group with the insulated bill at temperatures of 30°C (P = 0.0076) and 35°C (P <0.0001). At 35°C, the mean MR of the individuals with the insulated bills was 1.20 ml O_2_·g^-1^·h^-1^·°C^-1^ (± 0.06) while the intact group presented MR equal to 0.97 ml O_2_·g^-1^·h^-1^·°C^-1^ (± 0.07) (Figure 2). At ambient temperatures below 30°C, MR did not differ between treatments. The basal metabolic rate, measured within the TNZ, was equal to 0.779 ml O_2_·g^-1^·h^-1^·°C^-1^ (± 0.04) in the intact group and 0.733 ml O_2_·g^-1^·h^-1^·°C^-1^ (± 0.05) in the bill insulated group (*F*_1,6_=0.004; *P*=0.99).

**Table 1.**
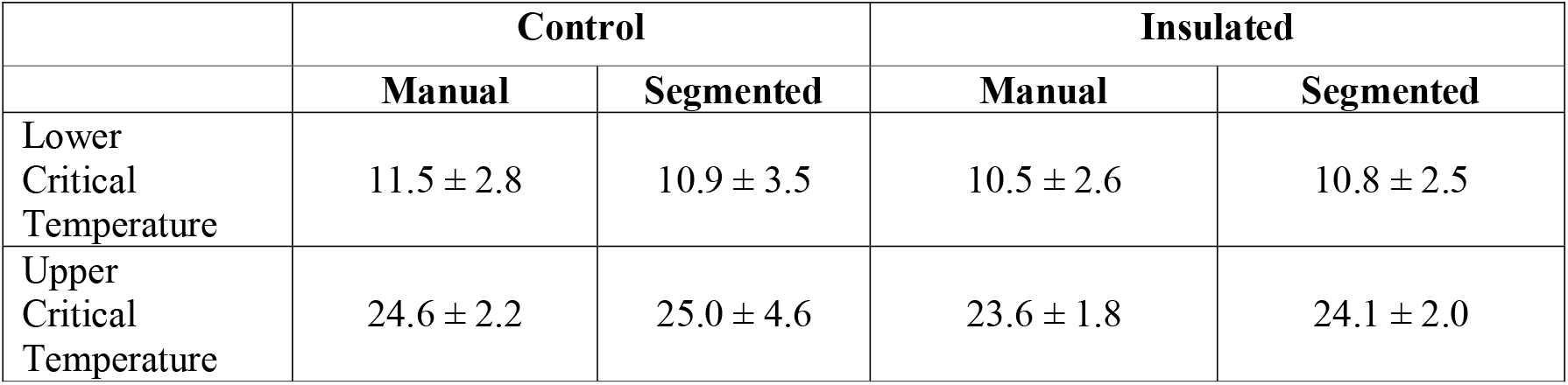
Lower and upper critical temperatures for the control group (intact bill) and experimental group (insulated bill) determined from the mass specific 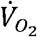 data using a manual intersection method contrasted with segmented regression approaches.

**Figure 2.**
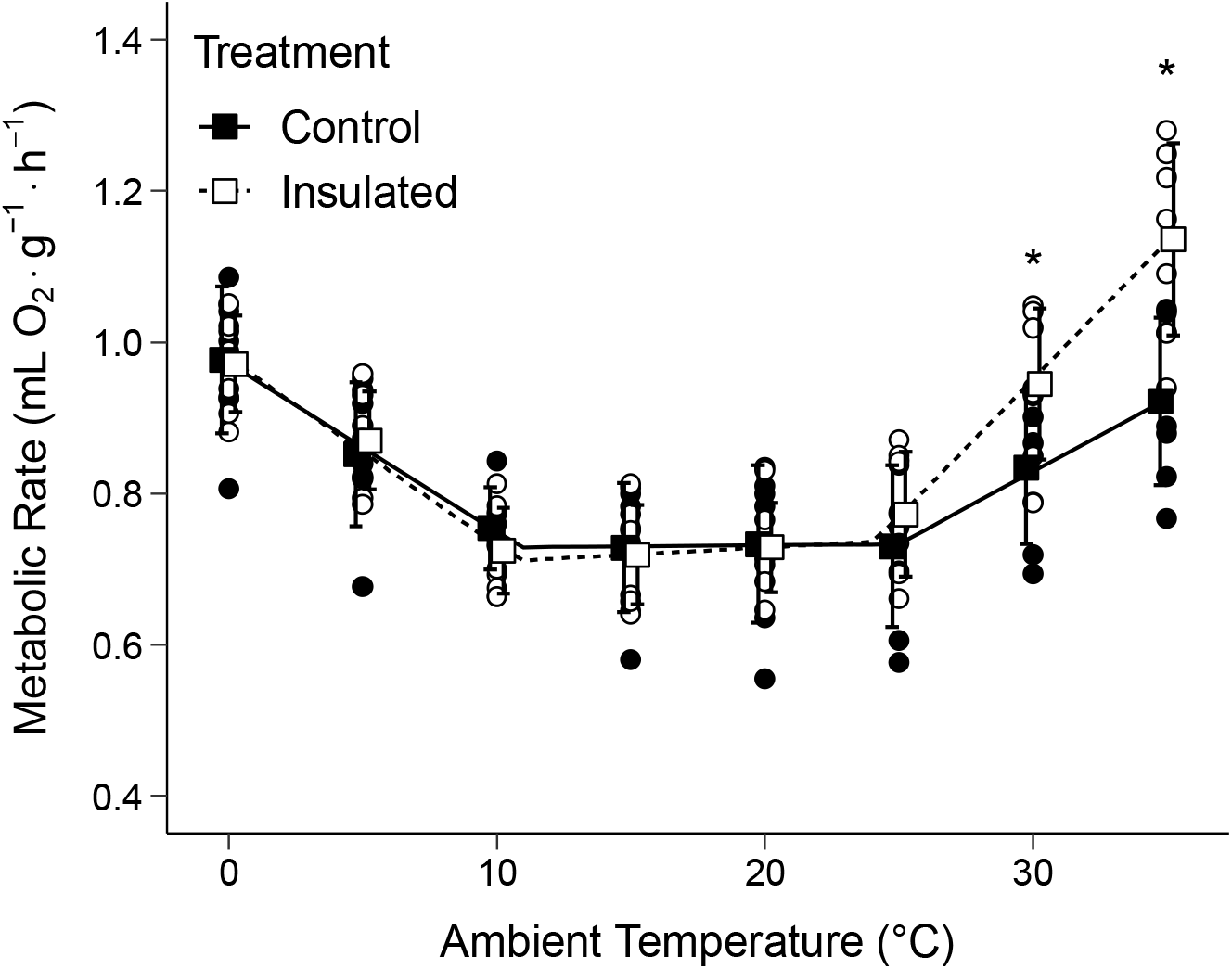
Metabolic rate of the toco toucan (*Ramphastos toco*) as a function of ambient temperature measured with the bill intact and the bill insulated. The asterisk symbol (*) represent the temperatures at which the treatments differed statistically.

The body temperature as a function of the ambient temperature did not differ between treatments (Treatment x Temperature Interaction: *F*_7,77.9_=1.53, *P*=0.17), although it was significantly related to ambient temperature (*F*_7,78_=36.28, *P*<2×10^−16^) in both treatment groups showing a rise in T_b_ as ambient temperature rose, and higher overall in the insulated condition (main effect of treatment: *F*_1,78.2_=5.69, *P*=0.019; Figure 3). Within the TNZ, the bird’s mean T_b_ was equal to 38.0°C (± 0.3) in the intact group and 38.2°C (± 0.5) in the bill insulated group.

**Figure 3.**
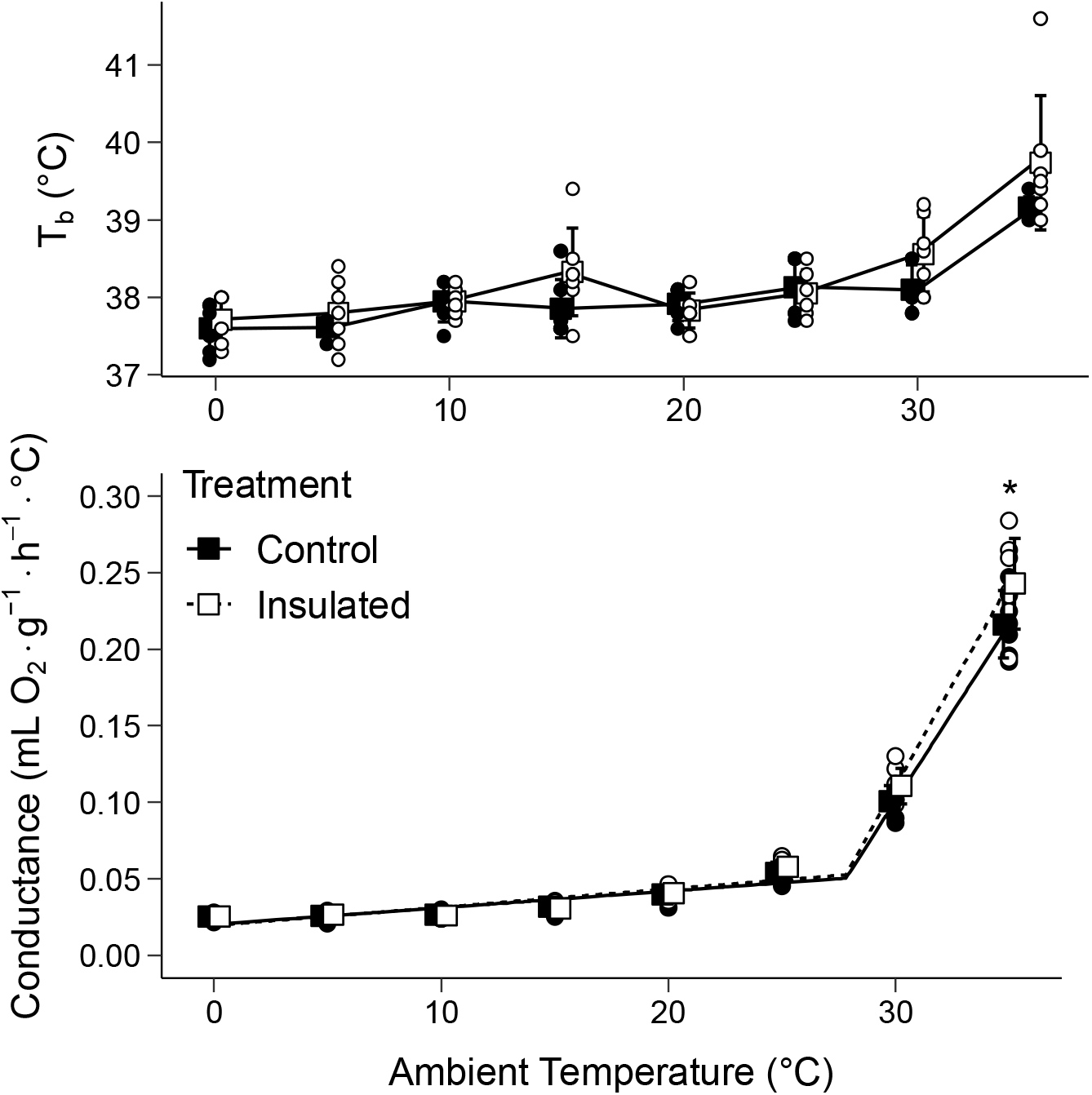
Body temperature (T_b_) (top panel) and thermal conductance (bottom panel) of toco toucan (*R. toco*) as a function of ambient temperature. Measurements were made with the bill intact (control) and with the bill artificially insulated. The asterisk symbol (*) represents the temperature at which the treatments differed statistically.

The relationship between thermal conductance (C) and ambient temperature could be described by a two-phase linear regression, intersecting at approximately 28°C for both intact and bill insulated birds (Figure 3; Table 2); this threshold was not affected by treatment (*F*_1,10_= 0.57; *P*=0.47). Overall, conductance was affected by temperature (*F*_7,79.3_ = 719; *P* < 2×10^−16^), by treatment (*F*_1,79.7_ = 7.77; *P*=0.0066), and by a temperature x treatment interaction (*F*_7,79.2_ = 719; *P*=0.0038). There was a marginally significant difference in C values at 30°C (*P*=0.05) between intact and bill insulated birds that became more pronounced at 35°C (*P*<0.001), at which point C was 1.16 times higher in the insulated group compared to the intact one. At ambient temperatures between 0 and 15°C, thermal conductance was minimal and virtually identical between both treatments, with mean values of 0.0276 (± 0.003) in the intact group and 0.0271 (± 0.002) in the bill insulated group.

**Table 2.** Upper threshold temperature for the control group (intact bill) and experimental group (insulated bill) determined from the mass specific conductance data using a manual intersection method contrasted with segmented regression approaches.

|  | **Control** |  | **Insulated** |  |
| --- | --- | --- | --- | --- |
|  | **Manual** | **Segmented** | **Manual** | **Segmented** |
| Upper<br>Threshold<br>Temperature | $27.8 \pm 0.4$ | $27.8 \pm 0.3$ | $28.0 \pm 0.5$ | $27.8 \pm 0.3$ |

Mean HP within the TNZ (i.e., BMR) was 2.66 ± 0.41 and 2.51 ± 0.21 in the control and bill insulated condition, respectively. Energetic costs, assessed from the difference in HP between insulated and control condition were significantly influenced by temperature (Figure 4; *F*_7,34.6_=9.01; *P*=2.7×10^−6^) and this effect was mostly caused by the large HP differential at the highest temperatures tested (i.e., 30 and 35°C, see Figure 4) since, in general, it hovered near 0 Watts for all other temperatures. At 35°C, for example, the cost of managing heat exchange in the bill insulated group was 0.797 W above the control group, which represented ∼31.7% of BMR; if this additional energetic cost was entirely offset by compensatory evaporative cooling mechanisms, it would correspond to the evaporation of 30.5 g H_2_O·day^-1^.

**Figure 4.**
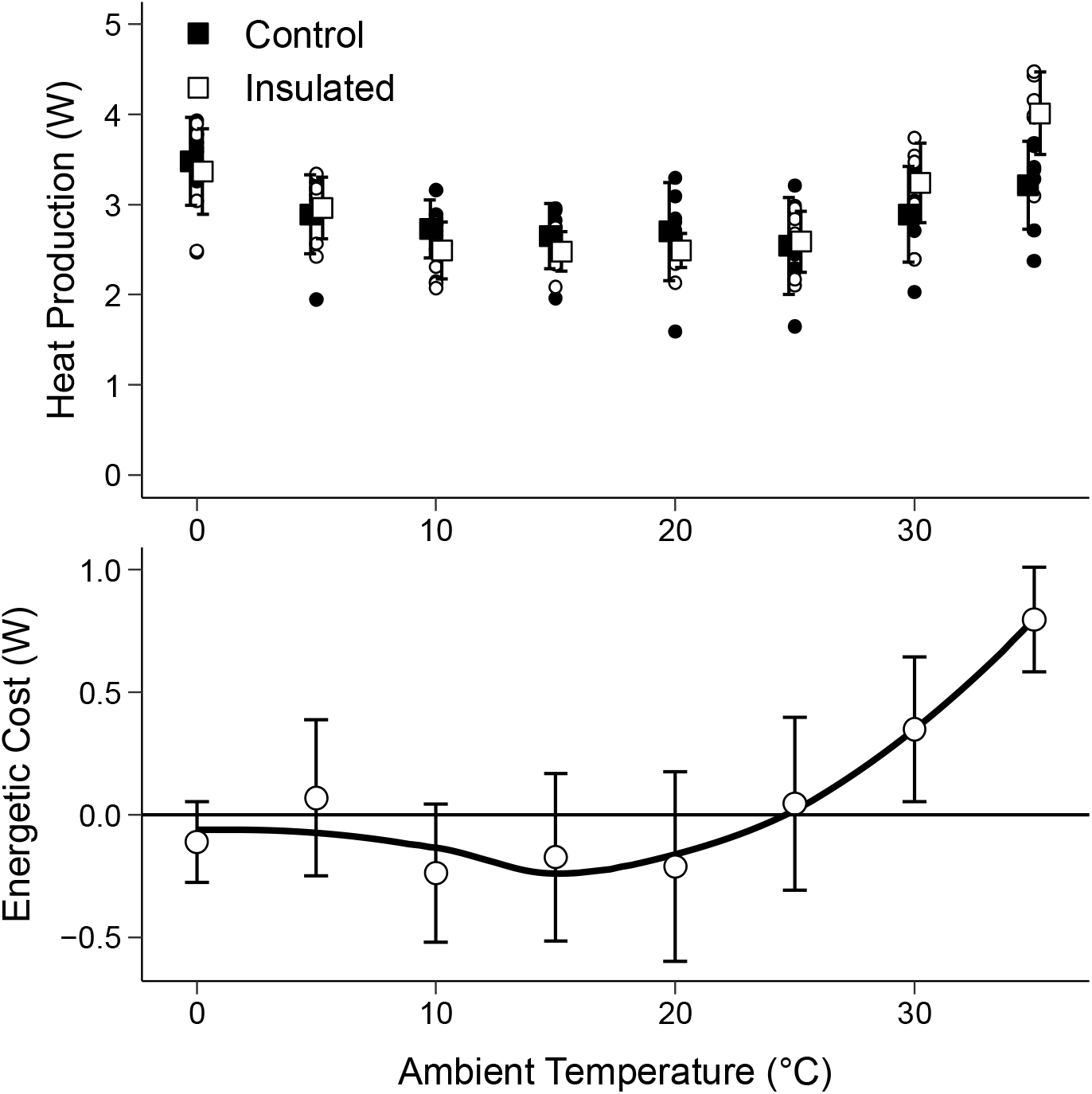
Heat production (HP) (top panel) and apparent energetic cost associated with insulating the bill (bottom panel) of toco toucan (*R. toco*) as a function of ambient temperature. Top panel depicts mean (± sd, with raw data points included) from both intact birds (control) and birds that had their bills covered (insulated). In the bottom panel, energetic cost (depicted by the open symbols) represents the difference between the respective (pairwise) measurements conducted during the insulated trial minus the control trial. Solid line depicts a simple smooth curve to aid in visualising the rise in energetic costs at higher temperatures.

## Discussion

Aside from the obvious role it plays in food acquisition, the avian bill’s role as a thermoregulatory organ has been well established for a diverse assemblage of birds subjected to different thermal habitat constraints (reviewed in Tattersall et al., 2017). Indeed, broad scale patterns correlating bill size and ambient temperature (McQueen et al., 2022; Ryding et al., 2021) provide further evidence that heat exchange considerations are of adaptive relevance and have played a role in the evolution of the anatomical attributes of this multi-task appendage (Tattersall et al., 2017). The bill assists in the maintenance of thermal homeostasis by efficiently modulating dry heat exchange via vasomotor adjustments as ambient temperature varies (Hagan and Heath, 1980; Tattersall et al., 2009). An intuitive and underappreciated consequence of this response, which causes an undetectable increment in metabolic rate, is that it alleviates the energetic cost of body temperature regulation (Gordon, 2012; McNab, 1974). Thus, one could expect that the greater the capacity of a bird to exchange heat through the bill, a trait that would co-vary bill size (e.g., surface area), the larger would be the range of ambient temperatures the animal is capable of withstanding without recruiting more energetically expensive thermoregulatory responses (i.e., the wider would be its TNZ). The results we obtained in the toco toucan clearly revealed that this was not the case, as we found no difference in the TNZ width between intact birds able to explore their extraordinary capacity for heat exchange through the bill at fullest extent and those where such a capacity was limited. Although contrary to our primary hypothesis, the closer examination of the patterns of metabolic variation at the full range of temperatures tested may yield some insights that we will cover in detail below.

The extreme capacity of toco toucans to dissipate heat through their bills provides an important advantage in assisting the birds in maintaining thermal homeostasis under high ambient temperatures or under situations in which heat production is increased (Tattersall et al., 2009). However, at the other extreme, at low temperatures, the possession of a large uninsulated appendage may represent a potential liability to heat conservation since underlying the most external bill layer, the ramphotheca, is composed of living tissue (Midtgard, 1984a; Midtgard, 1984b; Van Hemert et al., 2012) that will require some moderate level of circulatory irrigation even if dramatically reduced by a thermoregulatory-triggered vasoconstrictive response (e.g., Johansen and Bech, 1983). Therefore, we were expecting that intact toucans would lose more heat to the environment than the bill insulated ones under low temperature conditions and that this would cause metabolic rate to increase at a higher LCT in the intact condition. We found, however, that there was virtually no difference in LCT and birds in both treatments start to exhibit an increase in metabolic rate as temperature drops below ∼10.5-11.5°C. This observation points out that toco toucans are indeed able to finely tune heat dissipation through the bill and overall heat production to keep T_b_ constant at low temperatures, a response that was only previously implied (Tattersall et al., 2009). It also revealed how effective the vasomotor response of the toco toucan bill could be in promoting heat conservation at cold temperatures. Infrared imaging, however, shows that toucans keep a temperature differential of approximately 1 and 4°C between their distal and proximal bill surface, respectively, and ambient temperature, within the 10 to 20°C range (Tattersall et al., 2009). Therefore, some heat must be lost through the bill at low temperatures and the reason why this does not lead to differences in metabolic rate when the bill is insulated may rely on the fact that toco toucans combine their autonomic vasomotor thermoregulatory response with behaviour. When sleeping, toucans generally place the bill under one of the wings and raise the tail in order to cover the bill (Alvarenga, 2004). Indeed, in virtually all trials run below 10°C and whenever asleep below 25°C, we found the birds at the end of the experiment with the bill tucked between their wings, a widespread response that has been shown to mediate thermoregulation in many bird species (Pavlovic et al., 2019), particularly in those with larger bills (Ryeland et al., 2017). Indeed, the experimental bill insulation we devised had a thermal conductivity comparable to that provided by avian feathers (Rogalla et al., 2022; Walsberg, 1988; Wolf and Walsberg, 2000), therefore, we can conclude that the vasomotor and behavioural responses were as effective at heat conservation as having the bill surface entirely insulated, as tested in the present study.

Toco toucans start to exhibit a bill vasodilatory response at temperature around 20°C, which becomes prominent throughout the entire bill surface as it rises above 25°C (Tattersall et al., 2009). The temperature of 25°C agrees well with the UCT values we found for toucans, both in intact and in bill insulated birds. Also, we found no difference in body temperature between treatments, therefore, discounting any potential heat accumulation in the insulated bill group. Accordingly, birds with insulated bills must have recruited other heat dissipating mechanisms to compensate for the disablement of their main thermal window, the bill, and maintained thermal balance within the TNZ. These compensatory responses may have included a vasodilator response to the feet and legs (Chaves, Tattersall, and Andrade, unpublished data) and the modulation of feather’s fluffing (e.g., Downs and Ward, 1997; Saarela et al., 1984; Weathers et al., 2001). Unfortunately, we were unable to collect data on the rates of evaporative water loss during the metabolic measurements and, therefore, we cannot ascertain whether evaporative cooling had played a significant role in the regulation of body temperature within the TNZ limits. If that was the case, we suspect that it may have been restricted to a moderate increase in respiratory frequency (Tattersall et al., 2009), and, perhaps, to gaping (Neumann, 2016) rather than active panting as it did not translate into a noticeable increment in metabolic rate. Indeed, it has been demonstrated that the increment in evaporative cooling is often dissociated from the metabolically determined UCT (Gordon, 2012; Mitchell et al., 2018).

At the two temperatures tested above the UCT, 30 and 35°C, we found that bill-insulated birds had their metabolic rate significantly increased above those with intact bills. Again, there was no difference in body temperature between treatments and, therefore, it seems likely that insulated birds were heavily panting at these temperatures (Tattersall et al., 2009; van de Ven et al., 2016) which may explain their higher metabolic rate. We did observe, however, at elevated ambient temperatures (above the UCT) that the thermal conductance was higher in the bill insulated condition compared to the control (Figures 2 and 3), indicative of higher metabolic costs of thermoregulation under heat stress situations. Indeed, insulating the bill impacts the energetics of thermoregulatory responses to heat stress, presumably because bill insulated birds are forced to recruit compensatory thermoregulatory mechanisms to augment thermal conductance at a greater intensity compared to birds in the control group in which the thermoregulatory role of the bill was kept functional. We have not quantified these mechanisms, but they are likely to include changes in posture, behaviour, and alternative thermal window recruitment, and increased levels of evaporative water loss from panting or gular fluttering. Thus, having access to the bill as a thermoregulatory organ at higher temperatures might have two important implications, one energetic and the other related to water economy. In terms of energetics, our estimate is that the beneficial consequence of being able to access the normal heat exchange through the bill at 35°C corresponds to ∼32% of BMR. The second consequence of disabling the capacity for heat exchange through the bill at higher temperatures is that toucans are forced to resort to higher levels of evaporative cooling mechanisms to defend body temperature. Also at 35°C, we estimated that this hydroregulatory cost would be of ∼30 g H_2_O/day, comparing favourably to allometric predictions (Gavrilov and Gavrilov, 2019). Thus, we demonstrated formally that a peripheral heat exchanger is quite effective at minimising both energetic and hydroregulatory costs of thermoregulation, reaffirming previous suggestions that bill heat exchange can offset evaporative cooling in birds (Greenberg et al., 2012).

The importance of the toucan’s bill in controlling body temperature can also be appreciated from their ecology. The toco toucan has a Neotropical distribution, occupying open fields to high tropical forests (Short and M., 2002). In Brazil, large populations are found in Cerrado vegetation (Sick and Barruel, 1988), where high temperatures (40 - 41 ° C) are recorded during spring and summer (Alves and Rosa, 2008; Silva et al., 2008). In these environments, it is possible that the bill of the toco toucan contributes in an important way both to the maintenance of thermal homeostasis (Tattersall et al., 2009), energetic and water balance (present study). Indeed, the size of the birds’ thermal windows, particularly the bill, may be under selection in relation to the thermal environment they occupy (Larson et al., 2018; Symonds and Tattersall, 2010). While our measurements in the lab and at rest did not show any profound influence of bill heat exchange on the limits of the TNZ, under more natural conditions (e.g., activity, solar heat exposure), the importance of this heat exchange organ is likely to be much greater. For example, the performance of the toucan’s bill as an efficient thermal window could allow this species to remain active for longer during the hot hours of the day, dedicating this extra time to ecologically relevant activities, such as defense of territory, foraging, and reproduction (Luther and Danner, 2016; van de Ven et al., 2019). If that is the case, toco toucans may have an adaptive edge over other birds that occupy the same habitat and are of similar size but have smaller thermal windows.

In conclusion, the incredible vasomotor control over the bill blood supply exhibited by the toco toucan suggests that they can simultaneously have an effective radiator of body heat at high temperatures without paying an obvious cost at low temperatures. In other words, the fact that we found no difference in thermal conductance, metabolic rate, or body temperature from bill insulated and bill uninsulated birds clearly shows that the bill’s cold-induced vasoconstriction was as good at preventing heat loss as having the bill covered with a highly insulating material. Although we did not find differences in the limits of TNZ associated with insulation of the bill, we did find a very clear effect at the highest temperatures tested. At temperatures above UCT, the access to the bill as a heat exchange avenue promoted a lower energetic cost and a considerable water economy while birds were defending body temperature. These results corroborate the idea that thermoregulatory considerations might be of adaptive value in birds, especially for large billed species, such as the toco toucan. We did report an elevation in thermal conductance of the insulated group, suggesting compensatory changes in behaviour or the autonomic function of other thermolytic heat exchangers in the body. We also observed a large difference in energetics of insulated birds at the temperatures above the UCT. This latter observation suggests that toucans can almost fully compensate for the absence of bill heat exchange by adopting more costly thermolytic mechanisms, such as evaporative water loss through gaping and panting.

## Acknowledgements

We wish to acknowledge Guillherme Gomes and Ariovaldo Pereira da Cruz-Neto for assistance with experiments and preliminary data analysis, and Luá T. Timpone and Adriana Fuga for assistance with animal care.

## Data Availability

http://hdl.handle.net/10464/16884

## Author Contributions

Conceptualization: DVA, JNC, & GJT; Writing – Original Draft Preparation: JNC, GJT, & DVA; Writing – Review & Editing: GJT, DVA, & JNC; Investigation: JNC & DVA; Data Curation: GJT; Formal Analysis: JNC & GJT; Visualization: GJT; Funding Acquisition: DVA.

## Supplementary Material

### Thermal conductivity of the insulating material

To calculate the thermal conductivity (*k*) of the material that insulated the toucans’ beak, we used a copper plate (12.8 cm wide x 17.5 cm long x 0.3 cm thick) with two resistors in parallel (12 ohms) inside, which were used to control the temperature of the plate. This system was developed in the Neurobiophysics and Electronics laboratories of the IFSC at USP in São Carlos. To control the temperature of this system (T_s_), we used a DC power supply (model MPL – 1303) and we chose to generate a power of 7.8W, through a voltage of 20V and an electric current of 0.39A. At the ambient temperature tested (5°C), the T_s_ of the plate was kept close to the body temperature of the toucans (38°C). For all the other three temperatures tested (15, 25 and 35°C) we kept the same power (7.8W) to verify if there was a variation of k as a function of the ambient temperature.

Three temperature sensors (K-type thermocouples connected to a 4-channel RDXL4SD-OMEGA Systems data logger) were inserted into the surface of the copper plate, and the average value recorded between the three sensors was used to define the temperature. surface (T_s_) of the plate. The plate was then coated with the same insulating material used to cover the bill, and the edges were supported on 2 cm Styrofoam plates. The surface temperature (T_s_) of the felt was collected through photos taken by a thermographic camera (FLIR SC-640) and through a specific software (Thermocam Researcher Pro 2.9) we obtained, through analysis routines, a surface temperature of 6 bounded regions (50×100 pixels: ∼5000 surface temperature points per region). Finally, the felt T_s_ was defined by the average of the T_s_ of these six regions. A thermocouple also recorded the surface temperature of the insulator to verify those of the thermographic image.

The experimental apparatus was introduced into a climatic chamber measuring 3×4m (Eletrolab) to control the ambient temperature. The ambient temperature (T_a_) and relative humidity (RH) data were collected by a humidity and temperature sensor and recorder (model RHT10 Extech®). To verify if there was a relationship between the *k* values and the Ta/RH data, we used Pearson’s linear correlation coefficient.

For each temperature, the experiment lasted 40 minutes (time interval necessary for equilibrium to occur in the system). The average temperature of the chamber and the surface of the plate were calculated in the last 20 minutes, whereas the temperature of the felt surface was obtained right after the defined time interval (40 min). At that moment it was necessary to enter the room to obtain thermal images of both sides of the plate, to make sure that the T_s_did not change from one side to the other. Through these values it was possible to calculate the thermal conductivity (W·m^-1^·K^-1^) of the insulator from the following formula:

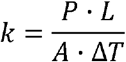

where P is the power (watts) supplied by the equipment, L is the thickness of the insulating material used, that is, the distance (meters) between the surface of the board and the surface of the felt, A is the surface area of the board (m^2^) and ΔT the temperature difference (Kelvin) between the surface of the plate and the surface of the felt.

## Results

The thermal conductivity (*k*) of the bill insulating material was positively related to the ambient temperature (*r* = 0.987; P = 0.013) and assumed an average value of 0.0729 (±0.006) W·m^-1^·K^-1^ among the four tested temperatures (Supplementary Table 1). The variation in the relative humidity (RH) inside the climatic chamber did not influence the *k* values (*r* = -0.649; P = 0.351).

**Supplementary Table 1.**
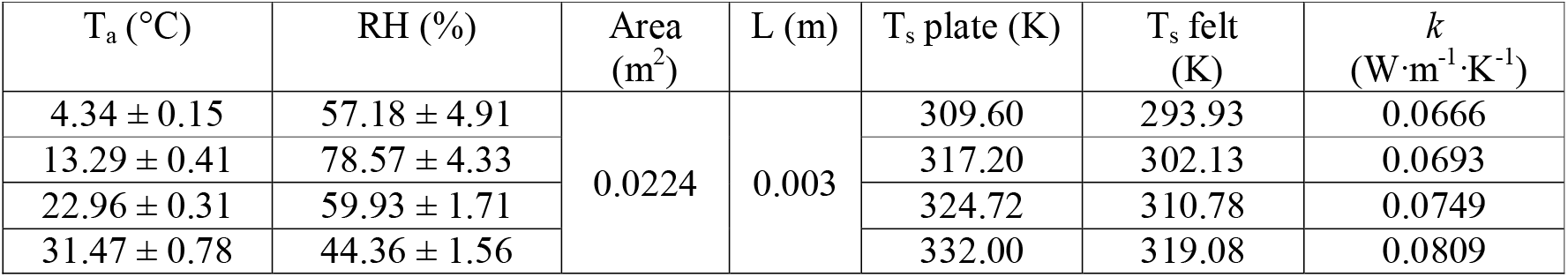
Mean values of ambient temperature (Ta) and relative humidity (RH) inside the climatic chamber and the variables used to calculate the thermal conductivity (k) of the insulating material of the bill. T_s_ = surface temperature.

